# Coadaptation of the chemosensory system with voluntary exercise behavior in mice

**DOI:** 10.1101/2020.05.24.113506

**Authors:** Quynh Anh Thi Nguyen, David Hillis, Sayako Katada, Timothy Harris, Crystal Pontrello, Theodore Garland, Sachiko Haga-Yamanaka

## Abstract

Ethologically relevant chemical senses and behavioral habits are likely to coadapt in response to selection. As olfaction is involved in intrinsically motivated behaviors in mice, we hypothesized that selective breeding for a voluntary behavior would enable us to identify novel roles of the chemosensory system. Voluntary wheel running (VWR) is an intrinsically motivated and naturally rewarding behavior, and even wild mice run on a wheel placed in nature. We have established 4 independent, artificially evolved mouse lines by selectively breeding individuals showing high VWR activity (High Runners; HRs), together with 4 non-selected Control lines, over 88 generations. We found that several sensory receptors in specific receptor clusters were differentially expressed between the vomeronasal organ (VNO) of HRs and Controls. Moreover, one of those clusters contains multiple single-nucleotide polymorphism loci for which the allele frequencies were significantly divergent between the HR and Control lines, i.e., loci that were affected by the selective breeding protocol. These results indicate that the VNO has become genetically differentiated between HR and Control lines during the selective breeding process, strongly suggesting the chemosensory receptors as quantitative trait loci (QTL) for voluntary exercise in mice. We propose that olfaction may play an important role in motivation for voluntary exercise in mammals.

## Introduction

Chemical senses are involved in many aspects of behavior. Olfaction is especially important for controlling such intrinsically motivated behaviors as food-seeking, social interactions, and reproductive- and fear-driven behaviors [1]. An ethologically relevant cue is detected by chemosensory receptors expressed in the sensory organs, which activate a specific neural circuitry for behavioral motivation and induces an appropriate behavioral output in a specific context. Comparative functional studies involving insect model species proposed a model wherein changes in chemoreceptor identity and expression are sufficient to provoke changes in neural circuit activity and behavioral outputs [2]. Thus, ethologically relevant cues, receptors, neural circuitries, and behavioral habits are likely to evolve together (coadapt) in response to natural and sexual selection.

One olfactory organ, the vomeronasal organ (VNO), occurs in some amphibians, squamates, and some mammals, including rodents. The VNO is known to detect intraspecific signals known as pheromones that trigger behavioral and physiological changes in receivers [3]. Pheromones are non-volatile peptides and small molecular weight compounds that are excreted in such fluids as urine and tears. These molecules are taken up from the environment to the VNO by direct contact and activate the vomeronasal sensory neurons (VSNs) [4, 5]. Generally, each VSN expresses a member of the vomeronasal receptor families: *type 1 vomeronasal receptors* (*Vmn1rs*), *type 2 vomeronasal receptors* (*Vmn2rs*), and *formyl peptide receptors* (*Fprs*), with some exceptions [6–10]. The signals detected by these receptors in the VSNs are axonally sent to glomerular structures and synaptically transmitted to the postsynaptic neurons, also known as mitral-tufted cells, in the accessory olfactory bulb (AOB) [11, 12]. The signals are then processed in the amygdala and hypothalamus, which induce the animal’s instinctive behavioral responses and endocrinological changes [3, 13, 14].

Rapid evolution of the receptor genes is a pronounced feature of the vomeronasal system [15–28]. Different species of animals have divergent family members of vomeronasal receptor genes [20, 23–25, 29, 30]. Even within the *Mus musculus* (house mouse) species complex, variation in the coding sequence is frequently observed [15]. Moreover, the abundance of receptor genes expressed in the VNO varies even among different inbred mouse strains [31]. Distributions of single nucleotide polymorphisms (SNPs) observed in lab-derived strains are non-random, and correlated with vomeronasal receptor phylogeny as well as genomic clusters [15]. These observations led us to hypothesize that selective breeding for a behavior that is modulated by chemosensory signals would induce an alteration in genomic clusters of vomeronasal receptors that are potentially involved in the behavior.

Voluntary wheel running (VWR) is an intrinsically motivated behavior, and even wild mice run on a wheel placed in nature [32]. Notably, VWR is one of the most widely studied behaviors in laboratory rodents [33–35]. Individual differences in VWR are highly repeatable on a day-to-day basis, the trait is heritable within outbred populations of rodents, and genes and genomic regions associated with VWR are being identified [36]. Moreover, some of the underlying causes of variation in VWR have been elucidated, in terms of both motivation and ability for voluntary exercise [34, 37, 38]. Importantly, a previous study demonstrated that the presence of conspecific urine increased VWR activity level in adult wild-derived mice [39], suggesting that external chemosensory cues also have a modulatory role in VWR activity.

We have established 4 independent, artificially evolved mouse lines by selectively breeding individuals showing high VWR activity (High Runners; HRs), along with 4 independent, non-selected Control lines over 88 generations [40, 41]. Briefly, all 4 HR lines run ∼2.5–3.0-fold more revolutions per day as compared with the 4 Control lines [42, 43]. Studies of mice allowed access to clean wheels or those previously occupied by a different mouse revealed that HRs show higher sensitivity to previously-used wheels and display greater alteration in daily wheel running activities than the Controls [44]. This result suggests that selective breeding for high running activity was accompanied by altered sensitivity to other individuals, suggesting a potential coadaptation of the chemosensory system with voluntary wheel running.

In this study, we examined whether selective breeding for VWR has differentiated the vomeronasal receptor genes between HR and Control lines. We found that a repertoire of receptor genes was differentially expressed between the VNO of HR and Control lines, which resulted from reduction or increase of specific vomeronasal receptor-expressing cells in the VNO of HR lines. We also found that this gene expression change was partially due to the genetic alteration upon selective breeding for VWR, suggesting a relationship between high running activity and the function of the VNO in HR lines. Taken together, our results indicate vomeronasal receptors as QTL for voluntary exercise behavior in mice.

## Results

### Differential expression of chemosensory receptors in the VNO of HR and Control lines

To examine the impact of selective breeding for VWR activity on receptor gene expression in the VNO, we conducted transcriptome analysis of the VNO from HR and Control lines. For each of the 4 HR and 4 Control lines, total RNA samples were prepared, each consisting of the combined VNOs from 3 individual males (Fig 1A). After RNA sequencing, we identified 76 differentially expressed (DE) genes in the HR line group compared to the Control line group (Fig 1B). There are 13 chemosensory receptor genes in the DE gene set, and all of them belong to either the *Fpr*, *Vmn1r* or *Vmn2r* family of the vomeronasal receptor genes (Fig 1B, shown in red). Of the 13 DE receptor genes, the Reads Per Kilobase Million (RPKM) of *Fpr3*, *Vmn2r8*, *Vmn2r9*, *Vmn2r11*, *Vmn2r96*, *Vmn2r98*, *Vmn2r102* and *Vmn2r110* were significantly up-regulated, while *Vmn1r188, Vmn1r236, Vmn2r15, Vmn2r16* and *Vmn2r99* were significantly down-regulated in the VSNs of HR lines compared to Control lines (Fig 1C). The RPKM of *olfactory marker protein* (*OMP*), which is abundantly and exclusively expressed in all mature VSNs in the VNO [45], was not different between HR and Control lines (Fig 1D), indicating that receptor gene expression changes were not due to variation in VSN number. The log◻ fold change of normalized expression of the DE genes varied from −3.4 to 2.0 (Fig 1E). *Vmn2r11* and *Vmn2r16* showed the largest upregulation and downregulation, respectively. These results suggest that expression of the chemosensory receptor genes is differentially regulated in the VSNs between HR and Control lines.

**Figure 1.**
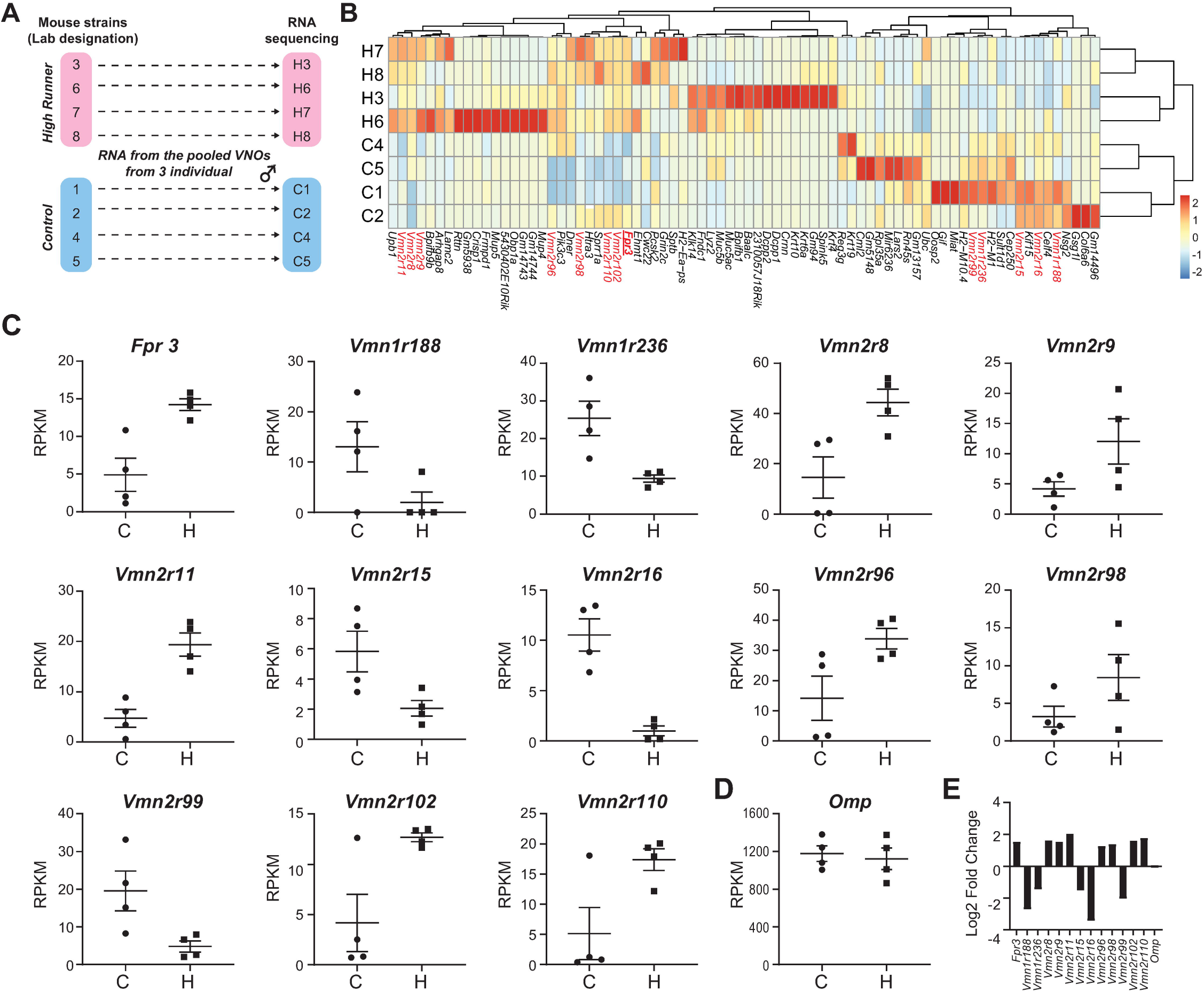
RNAseq analysis of the vomeronasal organs of High Runner and Control mice (A) Schematic of RNAseq sampling for analysis. (B) Heatmap of differentially expressed (DE) genes between HR and Control lines. DE vomeronasal receptors are shown in red. Fpr3 is highlighted with a red underline. (C and D) Scatter plots showing the RPKM of DE *vomeronasal receptor* (C) and *Omp* (D) genes in each line of HR or C mice. Error bars represent + S.E.M.. (E) Bar graph denoting log_2_ fold change of the relative expression of DE vomeronasal receptor genes between the HR and Control lines.

### Accumulation of all-or-none SNPs in a vomeronasal receptor cluster

We then hypothesized that differences between HR and Control lines in vomeronasal receptor gene expression would be associated with differences in allele frequencies between HR and Control lines caused by the selective breeding. Previous genome-wide SNP analysis detected 152 out of 25,318 variable SNP loci for which allele frequencies were significantly different between HR and Control lines after correction for multiple comparisons [46]. As explained in the previous paper, the differentiation in allele frequencies for these 152 loci cannot be attributed to random genetic drift. Of the 152 SNP loci, we particularly focused on 61 loci that were fixed for the same allele in all 4 replicate HR lines but not fixed in any of the 4 replicate Control lines, or vice versa (which we term “all-or-none SNPs”, Supplemental table 1). The 61 SNP loci were not randomly distributed throughout the genome (Supplemental Fig 1A). The majority of them (59 of 61) existed as a member of groups of 3 or more which were located in close proximity on the genomic chromosomes (Supplemental Fig 1B). As a result, only 11 all-or-none SNP clusters were observed in the genome (Supplemental Fig 1A)

Interestingly, 8 of the 61 all-or-none SNP loci were located in a ~3 Mb interval on chromosome 17 that contains clusters of *Vmn1rs* (14), *Vmn2rs* (21), and *Fprs* (7) (Fig 2A). Strikingly, 7 out of the 13 DE vomeronasal receptors are located in this all-or-none SNP cluster. Five of the all-or none SNPs are localized near the differentially expressed vomeronasal receptors (Fig 2B): SNP ID rs33447983 at 8.4 kb downstream of *Vmn2r99*, rs6224641 at 29 kb downstream of *Vmn2r99*, rs33649277 at 44 kb upstream of *Vmn2r102*, rs29522462 at 8.5 kb upstream of *Vmn2r109* and 524 bp downstream of *Vmn2r110*, and rs33120398 at 22 kb upstream of *Vmn2r109* and intron 1 of *Vmn2r110*. Other SNPs, such as rs33463529, are also closely located near vomeronasal receptors, though changes in expression of the nearby receptors were not observed. These results strongly suggest that changes in vomeronasal receptor gene expression between HR and Control lines are at least partially caused by changes in allele frequencies at multiple loci in response to selective breeding for VWR activity.

**Fig 2.**
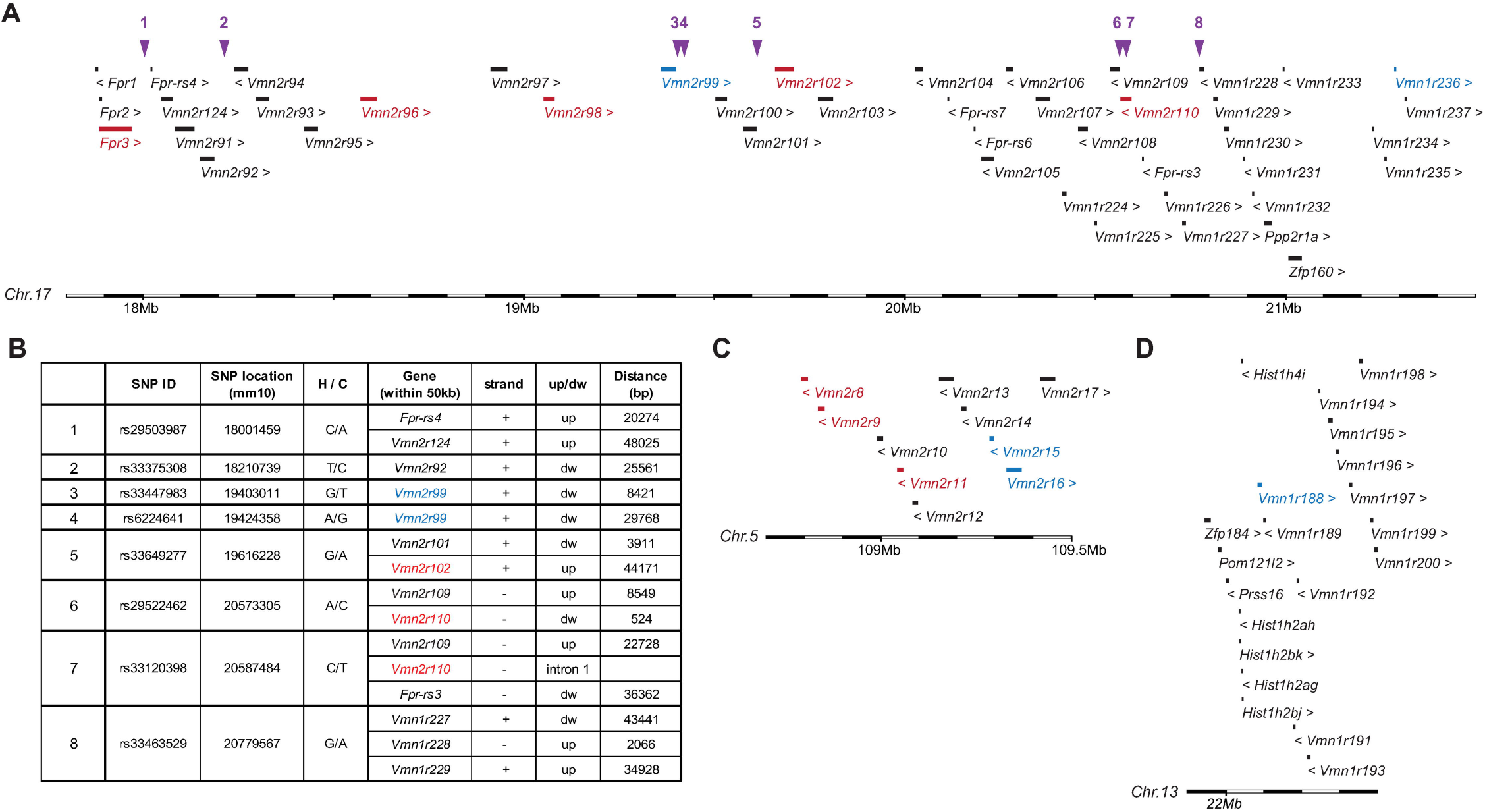
Genomic clusters containing the differentially expressed vomeronasal receptors (A, C and D) Genomic clusters of DE vomeronasal receptor genes in the mouse chromosomes 17 (A), 5 (C), and 13 (D). Vomeronasal receptors in red and blue indicate up- and down-regulations, respectively. Non-DE genes are shown in black. Purple arrowheads in (A) indicate locations of SNPs that are significantly differentiated between HR and Control groups [46], as shown in a table (B).

The rest of the DE vomeronasal receptors are located in another single ~1 Mb genomic cluster on chromosome 5 (Fig 2C), with one exception that is located on chromosome 13 (Fig 2D). Neither cluster contained SNP loci that are significantly differentiated between HR and Control lines [46]. Therefore, expression changes of the receptor genes in these clusters may be mediated by SNPs that remain polymorphic in both lines, or by different mechanisms.

### Differential number of Fpr3-expressing VSNs in the VNO of HR and Control lines

To determine the significance at the cellular level in the VNO of the DE chemosensory receptor genes, we chose one representative gene to determine whether there are differences in the number of receptor-expressing VSNs, or alternatively, differences in transcript abundance in each receptor-expressing VSN. We performed *in situ* hybridization to detect *Fpr3* using RNAscope *in situ* hybridization in the VNO of 2-3 individual mice from each of the 4 HR and 4 Control lines, together with a probe for the *Gαo* (*Gnao1*). Expression of *Fpr3* is ~3 times higher in HR lines compared to Control lines in RNAseq analysis (Fig 1C). Although *Fpr3*-expressing VSNs were observed in the VNO of both HR and Control lines, the number of *Fpr3*-expressing VSNs in each VNO slice varied among lines (Fig 3A, B). In 3 Control lines (line 1, 4, and 5), *Fpr3* signal was barely observed in each VNO tissue slice, while Control line 2 had a significantly higher number of *Fpr3*-expressing VSNs in each slice (Fig 3B, one-way ANOVA (*p* < 0.0001) with Tuckey’s *post hoc* test (*p* < 0.01)). This result was consistent with the RNAseq data, in which the amount of *Fpr3* transcripts in line 2 was higher than other Control lines (Fig 1B, highlighted with a red under line). On the other hand, we consistently observed multiple *Fpr3*-expressing VSNs in most of the VNO tissue slices from the 4 HR lines. The number of *Fpr3*-expressing VSNs in each HR lines was significantly higher than that of Control line 1, 4, and 5 (Fig 3B, one-way ANOVA (*p* < 0.0001) with Tuckey’s *post hoc* test (*p* < 0.05)) and did not differ among the 4 HR lines and Control line 2 (Fig 3B). Generally, there were significantly more *Fpr3*-expressing VSNs in the HR versus the Control lines (Fig 3C, unpaired t test (*p* < 0.05)), and the fluorescent intensity derived from *Fpr3* gene transcripts in each VSN was not distinguishable between the VNO tissues from HR and Control lines (Fig 3D and E). Taken together, these results demonstrate that the different expression levels of the chemosensory receptors result from changes in the number of receptor-expressing VSNs. Thus, the number of VSNs expressing specific sets of chemosensory receptors are differentially regulated after selective breeding for VWR.

**Fig 3.**
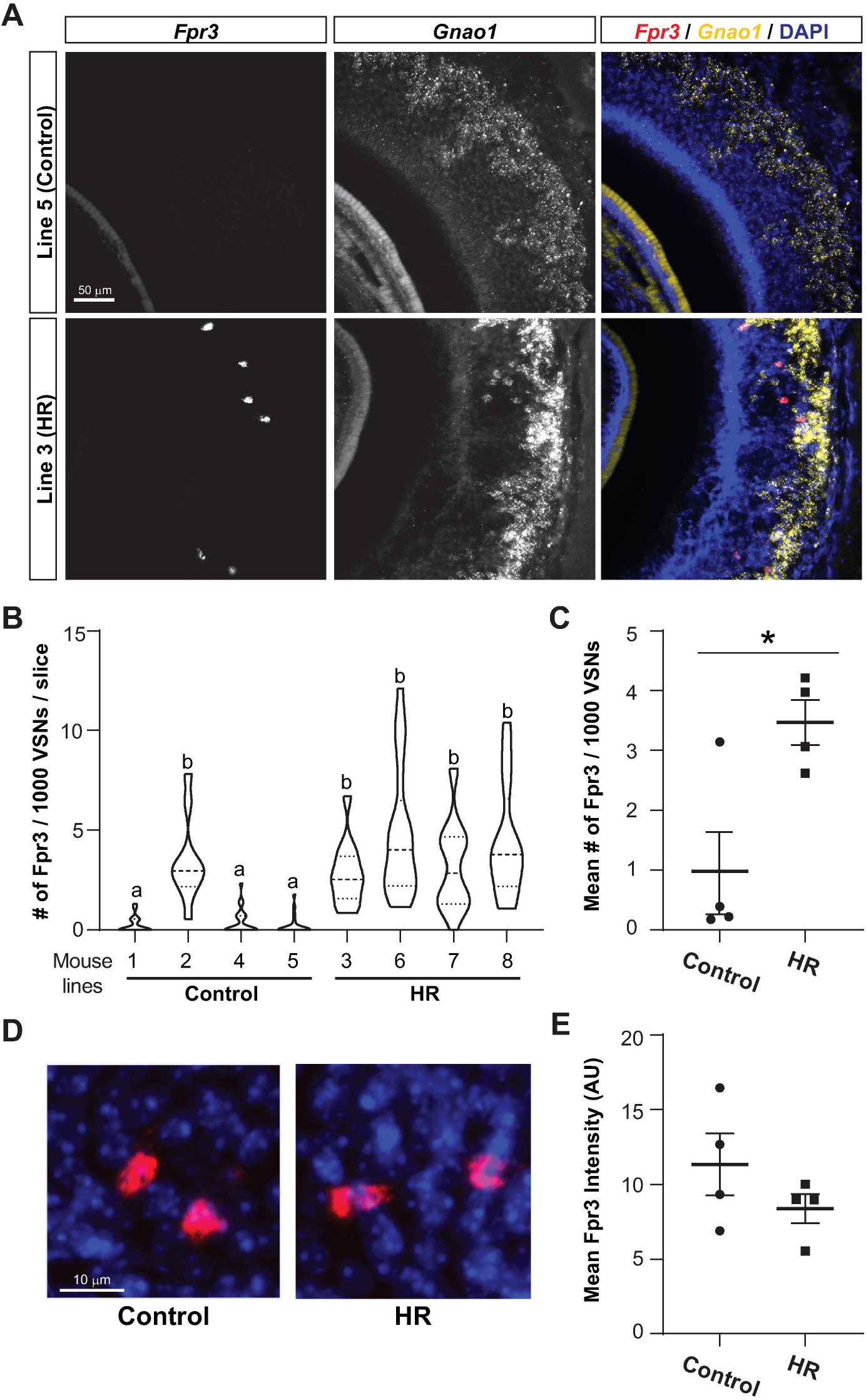
RNA scope *in situ* hybridization analysis of a DE receptor gene in the VNO (A) Images showing RNAscope-derived fluorescent signals for *Fpr3* (left) and *Gnao1* (middle) transcripts. In merged images (right), *Fpr3* and *Gnao1* are shown in red and yellow, respectively, together with DAPI staining (blue). Upper and lower panels show representative images from the VNO of a Control (line5) line and a HR (line3) line, respectively. (B) A violin plot showing the number of *Fpr3* signals in 1,000 vomeronasal sensory neurons per VNO slice for each line of mice. n = 10 - 22 slices in 2 - 3 mice per line. The plots with different letters are significantly different in ANOVA (*p* < 0.0001) with Tuckey’s *post hoc* test (*p* < 0.05). (C) The mean number of *Fpr3* signals in 10,00 VSNs in Control and HR lines. Each dot indicates the mean of one line. Error bars represent + S.E.M.. * indicates *p* < 0.05 in a *t*-test. (D) Representative images showing *Fpr3* signals (red) observed in the VNO of Control and HR lines of mice. DAPI signals are shown in blue. (E) The mean of *Fpr3* signal intensity (arbitrary unit, AU) per VSN in the Control and HR lines. Each dot indicates the mean of one line.

## Discussion

In this study, we utilized a unique animal model: 4 replicate mouse lines that have been experimentally evolved by selectively breeding individuals showing high VWR activity (HR lines), along with their 4 independent, non-selected Control lines maintained over 88 generations [40]. The HR and Control lines provide a strong model for determining the contribution of genetics to voluntary-exercise related traits [41]. In addition to the exercise ability-related genetic adaptations found after selective breeding [34, 41], several changes at the level of the central nervous system have also been identified, which contribute to elevation of VWR for HR mice [34, 37, 47]. Through SNP mapping analysis (Supplemental Fig 1), we found that 3 of the 61 all-or-none SNP loci that were fixed in all 4 replicate HR lines (but none of the 4 replicate Control lines) were located in a genomic cluster exclusively containing T-box genes on Chromosome 5. These genes are associated with GO terms of “bundle of His development,” “embryonic forelimb morphogenesis,” “cardiac septum morphogenesis,” “ventricular septum development,” and “cardiac muscle cell differentiation”. Indeed, compared with their 4 non-selected Control line counterparts, mice from the 4 replicate HR lines have been shown to have increased ventricular mass [42, 48–50], as well as altered cardiac functions [50–52]. Thus, the genome-wide SNP analysis of HR and Control lines of mice [46] could robustly identify QTL associated with voluntary exercise behavior.

Vomeronasal receptors are among the most rapidly evolving genes in vertebrates [15–28]. Different taxonomic groups have divergent family members of vomeronasal receptor genes [18, 20, 23–25, 29, 30], and the abundance of receptor genes expressed in the VNO is different even among inbred mouse strains [31]. Moreover, many of the mouse pheromones identified as ligands for vomeronasal receptors show strain specificity. For example, expression of the male pheromone ESP1 is only observed in a few inbred strains, although males of wild-derived strains all secrete abundant ESP1 peptide into their tears [53]. Likewise, expression of juvenile pheromone ESP22 is missing in some inbred strains [54]. Major urinary proteins (MUPs) are potential ligands for vomeronasal receptors, and all male mice of a given inbred strain secrete identical MUP members, whereas wild-derived mice each exhibit a unique profile of emitted MUPs [55]. Thus, pheromones and vomeronasal receptors in the vomeronasal system may have evolved in response to various environmental changes, including domestication, which resulted in alteration of coding sequences and expression patterns.

Considering the extensive evolution of receptor genes, selective breeding for a chemosensory-mediated behavior is an attractive alternative approach to reveal the functions of vomeronasal receptors. VWR activity of a mouse strain that recently derived from the wild has been shown to be increased by urinary chemosignals from other individuals [39]. Therefore, if the function of the VNO is involved in the modulation of VWR activity, then we would expect that selective breeding for high VWR activity should impact vomeronasal receptors. Indeed, we found that expression levels of several vomeronasal receptor genes as well as a few SNPs near the DE receptor genes were different between HR and Control lines. Although the role of each DE receptor in VWR activity needs to be determined in future studies, the current results suggest that vomeronasal chemosensory receptors could be important QTLs for voluntary exercise in mice.

One of the important remaining questions is how the vomeronasal system modulates VWR behavior in HR lines. One study measuring patterns of brain activity using c-Fos immunoreactivity revealed multiple areas in the brain that appear to be associated with motivation for VWR in HR lines [56]. These areas include brain nuclei known to be motivation-related, such as the prefrontal cortex, medial frontal cortex, and nucleus accumbens (NAc) [56]. In addition, it was recently shown in mice that VNO-mediated signals regulate the mesolimbic dopaminergic system, especially by upregulating the ventral tegmental area (VTA)-NAc circuit, and that they enhance reproductive motivation in mice [12]. Thus, it is possible that the VNO-mediated chemosensory signals also upregulate VWR activity by stimulating the VTA-NAc circuit. Moreover, one of the hypothalamic targets of the vomeronasal system, the medial preoptic area (MPOA), has been shown to regulate wheel-running activity in a hormone-dependent manner [57–60]. It is therefore also conceivable that the VNO-mediated chemosensory signals upregulate VWR by directly activating MPOA neurons. Combined with these previous observations, we propose that chemosensory signals detected by the VNO activate specific areas of the central nervous system that contribute to VWR activity. Future studies are expected to reveal the role of the VNO in modulating physical exercise and other voluntary behaviors in rodents.

## Materials and Methods

### Animals

The experimental procedures were approved by the UCR Institutional Animal Care and Use Committee and were in accordance with the National Institutes of Health Guide for the Care and Use of Laboratory Animals. The VNOs studied were from 12-week old male and female mice of 4 lines selected for high voluntary wheel running and 4 Control lines. The studied mice were derived from generation 88 of a replicated selective breeding experiment for increased voluntary wheel running behavior on the Hsd:ICR strain [40]. Wheel revolutions were recorded in 1-minute intervals continuously for 6 days, and mice were selected within-family for the number of revolutions run on days 5 and 6. In each selected HR line, the highest-running male and female within 10 individual families were selected per generation and each mouse was mated to a mouse from another family, within its line. This within-family selection regimen minimized inbreeding such that the effective population size was approximately 35 in each line [40]. In the Control lines, one female and one male within each family were chosen at random, though full sibling mating was again prevented. The mice in the present study were neither full nor half-siblings.

### RNA sequencing

The VNO tissues were harvested from 3 male mice from each of the 4 HR and 4 Control lines, immediately transferred to RNA later (Sigma-Aldrich), then stored at −80°C until use for RNA-seq. VNO tissues from the same line of mice were pooled and homogenized in Trizol Reagent (Life Technologies, Carlsbad, CA) and processed according to the manufacturer’s protocol. Trizol-purified RNA samples were quantified using Qubit1 2.0 (Life Technologies). The integrity of isolated RNA was measured by the 28S/18S rRNA analysis using the Agilent 2100 Bioanalyzer (Agilent Technologies, Santa Clara CA) with RNA Nano chip (Agilent Technologies, Palo Alto, CA). Samples had RNA integrity number values of at least 8.30. Using the Ultra II Directional RNA Library Prep kit (NEB), each RNA sample was depleted of ribosomal RNA and used to prepare an RNA-seq library tagged with a unique barcode at the UCR IIGB Genomics Core. Libraries were evaluated and quantified using Agilent 2100 Bioanalyzer with High Sensitivity DNA chip, then sequenced with the Illumina NextSeq 500 system (Illumina, San Diego, CA, USA) and 75nt-long single-end reads were generated at the UCR IIGB Genomics Core. A total of 8 libraries (4 HR lines and 4 Control lines) were multiplexed and sequenced in a single lane which yielded ~11,000◻M reads, averaging ~1,400◻M reads per sample.

The RNA-seq data files are available in the National Center for Biotechnology Information Gene Expression Omnibus (GEO) database (accession identifier GSE146644).

### Differential gene expression analysis

The analysis compared the transcriptome profiles from the HR and Control lines of mice. Quality control of the sequence reads included a minimum average Phred score of 30 across all positions using FastQC. Sequencing reads were aligned to the mouse reference genome (GRCm38/mm10), using STAR aligner ver. 2.6.1d [61] with an increased stringency unforgiving any of mismatches per each read (‘-outFilterMismatchNmax 0’). Any reads that map to multiple locations in the genome are not counted (‘-outFilterMultimapNmax 1’) since they cannot be assigned to any gene unambiguously. In order to determine the differentially expressed (DE) genes, generated BAM files were accessed with Cuffdiff [62], a program included in Cufflinks. Cuffdiff reports reads per kp per million mapped reads (RPKM), log◻ fold change, together with *p*-value, and *q*-values. After Benjamini-Hochberg false discovery correction, genes with adjusted *p*-values less than 0.05 were considered as DE genes.

### Analysis of all-or-none SNPs

SNP data in supplemental table 7 (Data_S7) in Xu and Garland (2017) [46] were used for this analysis. SNPs that separate all 4 HR and 4 Control lines (which we term all-or-none SNPs) were selected (Supplemental Table 1) and mapped onto mouse genome (NCBI37/mm9) using UCSC Genome Browser (https://genome.ucsc.edu). We noticed that most of the all-or-none SNPs occurred in groups. Thus, we mapped those SNP clusters onto genomes (Supplemental Fig 1A). Each cluster was defined - and + 0.1 Mb from the first and last SNP, respectively, observed in a specific location of the genome. Information of coding genes in each SNP cluster were extracted (Supplemental Fig 1B), For some clusters, Gene-to-GO mappings was performed with PANTHER (http://pantherdb.org).

### RNAscope *in situ* hybridization

Female mice (11 Controls and 9 HRs) were utilized for this analysis. The animals were intracardially perfused with 4% Paraformaldehyde in Phosphate Buffered Saline (PBS). VNOs were dissected from perfused animals and fixed overnight. The VNO samples were decalcified in EDTA pH 8.0 for 48 hours, then cryoprotected in 15% sucrose in PBS followed by 30% sucrose in PBS. All samples were ultimately embedded in optimal cutting temperature (OCT) medium (Electron Microscopy Sciences) above liquid nitrogen and sectioned at 20 μm using Leica CM3050S Cryostat. Slides were stored at −80◻C until use for *in situ* hybridization staining.

RNA detection in VNOs were performed with ACD RNAscope® control and target GNAO1 (ACD # and FPR3 (ACD #503451) using RNAscope® Multiplex Fluorescent Reagent Kit v2 (ACD# 323100) Assay. Probe binding was detected with Akoya Biosciences’ Opal 690 (FP1497001KT) and 570 (FP1488001KT) Dyes at 1:750 dilution in RNAscope TSA Buffer. Nuclear staining was visualized with DAPI (EMS #17989-20). Images were acquired at 20X or 40X magnification on Zeiss Axio Imager.M2, and FPR3-positivity was quantified with a proprietary script using QuPath software. Fluorescent intensity was measured by Fiji software. 4-8 slices in each animal were examined. One-way ANOVA with Tuckey’s *post hoc* test was used in Fig 3B. An unpaired t-test was used to examine statistical significance in Fig 3C and E.

## Supporting information

SupplementalFigure1-Nguyen

SupplementalTable1-Nguyen

## Acknowledgments

We thank Drs C. R. Yu, H. Matsunami, T. Girke, P. Campbell, C. A. Scott, M. Riccomagno and N. Yamanaka for helpful discussion and comments on the manuscript. We are grateful for technical assistance from Office of Campus Veterinarian, and Genomics Core Facility at University of California, Riverside (UCR).

## Author contributions

S.H.Y. and T.G. designed the project. Q.A.N., D.H., and C.P. performed experiments. Q.A.N., S.K., T.H., T.G. and S.H.Y analyzed data. S.H.Y wrote the manuscript with comments from all authors.

---

Supplemental Table 1. Genomic locations, *p*-value by the mixed model approach [46], and allele frequencies of the 61 all-or-on SNP loci.

Supplemental Fig 1. Analysis of all-or-none SNP loci in HR and Control lines of mice (A) A schematic diagram showing the relative positions of loci containing 1 or more all-or-none SNPs. Blue triangles indicate non-chemosensory clusters, and a red triangle indicates clusters containing only chemosensory (vomeronasal) receptors. (B) A table showing chromosomal location and length of each all-or-non SNP cluster, and the number of SNPs and genes within the clusters. The row highlighted in red is the cluster containing only the vomeronasal receptor genes.

