## Supplementary figures and images for "Coadaptation of the chemosensory system with voluntary exercise behavior in mice"

### SupplementalFigure1-Nguyen

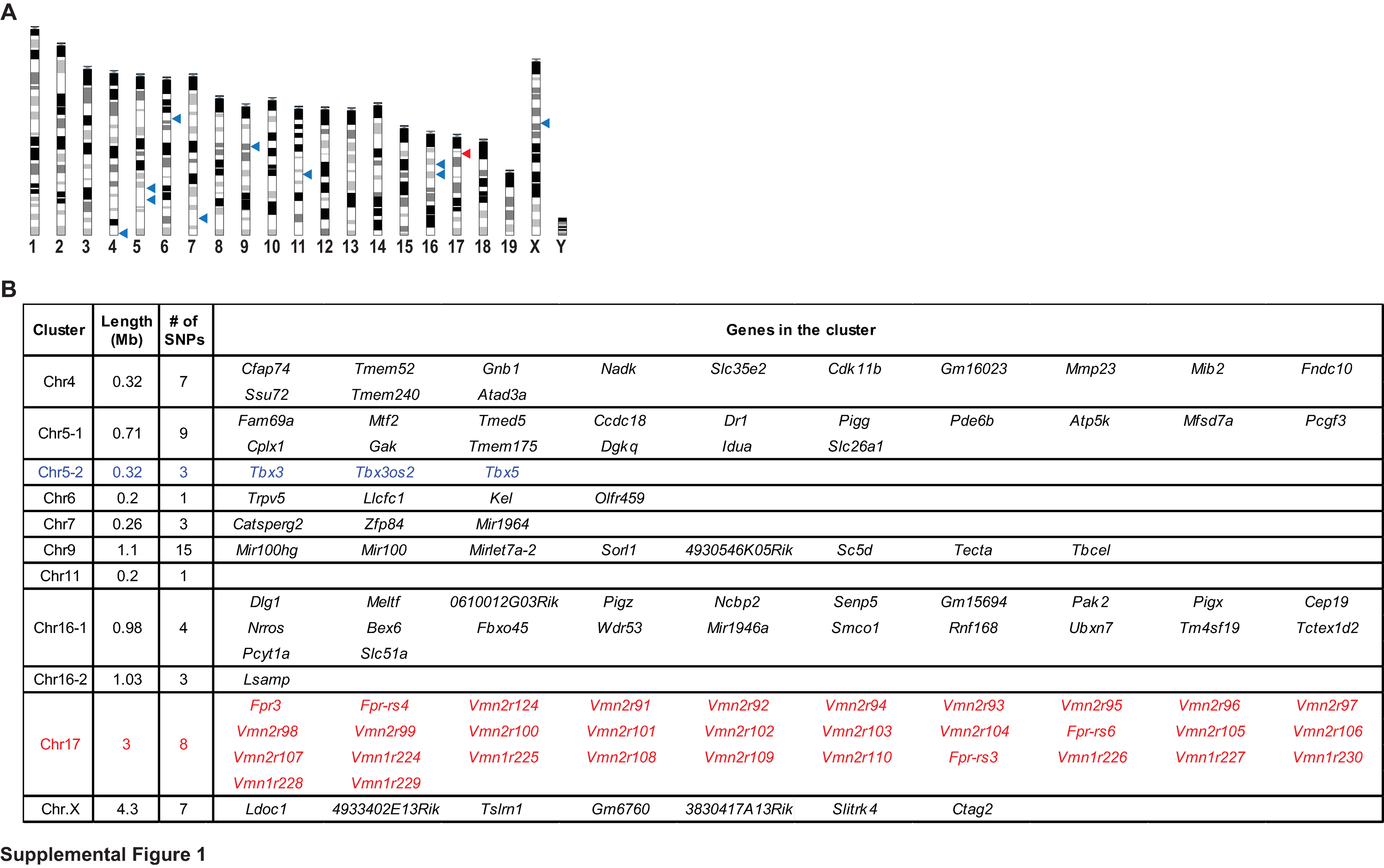
